# Distinct gene regulatory signatures of dominance rank and social bond strength in wild baboons

**DOI:** 10.1101/2021.05.31.446340

**Authors:** Jordan A. Anderson, Amanda J. Lea, Tawni N. Voyles, Mercy Y. Akinyi, Ruth Nyakundi, Lucy Ochola, Martin Omondi, Fred Nyundo, Yingying Zhang, Fernando A. Campos, Susan C. Alberts, Elizabeth A. Archie, Jenny Tung

## Abstract

The social environment is a major determinant of morbidity, mortality, and Darwinian fitness in social animals. Recent studies have begun to uncover the molecular processes associated with these relationships, but the degree to which they vary across different dimensions of the social environment remains unclear. Here, we draw on a long-term field study of wild baboons to compare the signatures of affiliative and competitive aspects of the social environment in white blood cell gene regulation, under both immune stimulated and non-stimulated conditions. We find that the effects of dominance rank on gene expression are directionally opposite in males versus females, such that high-ranking males resemble low-ranking females, and vice-versa. Among females, rank and social bond strength are both reflected in the activity of cellular metabolism and proliferation genes. However, pronounced rank-related differences in baseline immune gene activity are near-absent for social bond strength, while only bond strength predicts the fold-change response to immune (lipopolysaccharide) stimulation. Together, our results indicate that the directionality and magnitude of social effects on gene regulation depend on the aspect of the social environment under study. This heterogeneity may help explain why social environmental effects on health and longevity can also vary between measures.

## INTRODUCTION

Many animal species, including humans, live the majority of their lives as part of a larger group of conspecifics. Social group living provides a number of benefits, including protection from predators, improved territory and resource defense, and access to potential mates [1–4]. At the same time, it also generates competition for resources among group members. For many group-living species, the outcome of competitive interactions is at least partially predictable, giving rise to an observable social dominance hierarchy in which high status animals are consistently able to displace lower status animals [5–7]. Due to correlated differences in resource access, energy expenditure, and/or psychosocial stress, high-ranking and low-ranking animals are frequently behaviorally and physiologically distinct. For example, across social mammals, low status individuals often have elevated glucocorticoid levels or exhibit signs of glucocorticoid resistance [8–12].

However, correlations between social status and physiological measures are highly heterogeneous across species or between sexes, and sometimes even directionally inconsistent [8,13–18]. This heterogeneity is likely explained in part by differences in how status is attained and maintained. In some cases social status depends on individual characteristics, such as the ability to physically dominate competitors (e.g., male bottlenose dolphins, male red deer, female meerkats: [19–21]). In contrast, other types of social hierarchies are determined via nepotism, and do not strongly covary with individual phenotype (e.g., female spotted hyenas, some female cercopithecine primates: [22,23]). Hierarchies that are largely determined by physical condition are often dynamic, whereas nepotistic hierarchies can remain highly stable over time, and even extend across generations [24–26]. Consequently, while rank is an important predictor of fitness in both types of hierarchies [27–31], its physiological signatures may differ. For example, while high rank predicts lower glucocorticoid levels in female blue monkeys, female baboons, and naked mole-rats of both sexes [10,13,32,33], glucocorticoid levels tend to be higher in high rank female ring-tailed lemurs, female meerkats, and male chimpanzees [18,34,35].

In addition to the competitive interactions that structure social hierarchies, group-living animals can also form individually differentiated, affiliative social bonds. The affiliative behaviors that give rise to social bonds (e.g., proximity or contact in cetaceans and ungulates, grooming and proximity in primates) are often patterned, at least in part, by social status [36–42]. For example, in cooperatively breeding meerkats, dominant males groom dominant females more often than they groom subordinate females [43]. Similarly, attraction to high-ranking individuals commonly structures grooming patterns in social primates [40]. However, rank is not the sole determinant of affiliative behavior and social bond formation. In female yellow baboons, for instance, a measure of female social connectedness to other females is better predicted by the presence of maternal kin than by rank (although rank, not the presence of maternal kin, predicts female social connectedness to males; [44]). Recent evidence also indicates that the fitness effects of affiliative social relationships are also partially independent of rank. Stronger social relationships predict natural lifespan in members of at least five mammalian orders, and this relationship often persists after controlling for variation in rank or other measures of social status [38,45–51]. Indeed, in yellow baboons, social relationships predict lifespan even when rank does not [45].

Social status and social integration are therefore connected dimensions of the social environment that nevertheless can have distinct fitness consequences. This observation presents a puzzle about the mechanisms that make their consequences for health, physiology, and survival distinct. To date, far more work has focused on the physiological and molecular correlates of social status than of affiliative social bonds in natural animal populations. However, four lines of evidence argue that differences in affiliative social interactions should also be reflected in physiological or molecular variation. First, such changes are implied by cross-taxon support for an association between lifespan and social integration [52], suggesting at least a partial basis in physical condition. Second, studies in a small set of natural populations have already identified links between affiliative relationships and biomarkers of stress, especially glucocorticoid levels. For example, urinary glucocorticoids are lower in chimpanzees sampled while interacting with closely bonded social partners than in those interacting with non-bonded partners [53], and male rhesus macaques and female chacma baboons with stronger social bonds show reduced glucocorticoid responses to environmental stressors [54–56]. Third, social isolation and loneliness are associated with changes in human biology, including increased proinflammatory activity [57–59], hypothalamic-pituitary-adrenal axis activation [60,61], and risk for cardiovascular disease [62,63]. Finally, studies in captive rodents show that manipulation of social integration and social support can causally alter glucocorticoid regulation and increase the risk of cancer metastasis [64,65].

Despite these findings, most studies consider either the physiological signature of social status or of affiliative social relationships, not both. Further, those studies that incorporate both dimensions often measure only a single outcome variable in one type of social status hierarchy (i.e., physical competition-based or nepotistic). Because single measures vary along only one dimension, they have limited ability to distinguish shared versus unique signatures of competitive and affiliative interactions. Thus, it is possible that physiological changes in response to the social environment converge on a generalized signature of stress and adversity, in which low status and weak social bonds produce undifferentiable responses (e.g., the “conserved transcriptional response to adversity”: [66]). Alternatively, different facets of the social environment may be reflected in different biological pathways. If so, higher dimensional measures of physiological or molecular state may be informative about multiple aspects of an animal’s social experience, and help uncover why social status and social affiliation can be related, yet have distinct effects on fitness.

Functional genomic analyses of gene regulation provide an opportunity to differentiate these hypotheses. Importantly, previous work demonstrates the sensitivity of gene regulation to the social environment. For example, competitive interactions to establish dominance rapidly alter DNA methylation, histone marks, and gene expression across multiple vertebrate and social insect species [67–73]. Affiliative interactions can also be reflected in altered gene expression patterns. For example, experimental social isolation in piglets results in increased plasma cortisol and altered glucocorticoid and mineralocorticoid receptor expression in stress-related regions of the brain [74]. However, the species that have been central to understanding the genomic signatures of social status and social competition (e.g., cichlid fish, mice) tend not to be the same ones developed as models for social affiliation (e.g., voles, titi monkeys). Additionally, few studies of social interactions and gene regulation have focused on species that establish both clear social dominance hierarchies and long-term social bonds outside the mating pair-bond.

To begin addressing this gap, this study draws on data and samples from a five decade-long field study of wild baboons in Kenya, in which the fitness consequences of both social status and social relationships have been extensively investigated in prior work [29,44,45,75–78]. Gene regulatory signatures of the social environment have also been detected in this population [15,79]. Most relevant to this work, high-ranking baboon males exhibit elevated expression of inflammation-related genes both at baseline and upon stimulation with lipopolysaccharide (LPS; a pathogen-associated molecular pattern associated with gram-negative bacteria, and a strong driver of the innate inflammatory response: [15]). In contrast, little signature of rank was detectable in females [15]. This result is consistent with findings that high rank in males (but not females) predicts accelerated epigenetic aging, elevated glucocorticoid and testosterone levels, and, to a lesser extent, higher mortality rates [14,45,80]. However, it contrasts with the hypothesis of a highly consistent gene regulatory response to the social environment [66].

Together, these observations raise key questions about the extent to which the links between social experience and gene regulation are sex- and/or context-specific. To address them here, we expand on our previous white blood cell gene expression data sets by 64% (from n = 119 to n = 195 samples, including paired baseline and LPS-stimulated samples from nearly all individuals; Table S1). We also generated ATAC-seq data on chromatin accessibility in baseline and LPS-stimulated samples to infer the transcription factor binding events that explain social environment associations with gene expression. Our results indicate even more substantial sex differences in the signature of dominance rank than was apparent in previous work. We also identify, for the first time in any natural vertebrate population, a strong signal of social bond strength on gene regulation. Although several of the major pathways associated with rank and social bond strength overlap, their signatures are clearly distinct, and only social bond strength predicts the gene expression *response* to pathogen stimulation (i.e., the difference between baseline and LPS-stimulated cells from the same individual). Together, this work emphasizes the strong relationship between the social environment and gene regulation in the immune system in wild social mammals. It thus deepens our understanding of how fitness-relevant social experiences “get under the skin” to affect health and fitness outcomes.

## RESULTS

### Directionally opposite gene expression signatures of dominance rank in male and female baboons

We used RNA-seq to measure genome-wide gene expression levels in white blood cells from 97 unique adult baboons (45 females, 52 males; Table S1). For each animal, we collected RNA from paired baseline (unstimulated cells cultured in media) and LPS-stimulated samples that were cultured in parallel for 10 hours (Fig 1A; following [15]). Following quality control, our data set consisted of RNA-seq data from 195 samples (119 samples from previously published work and an additional 76 samples that are newly reported here; 97 unique individuals total, with 3 individuals represented by more than one blood draw; Table S1). Samples were sequenced to a mean coverage of 17.4 million reads ± 7.7 million s.d. (Table S1). After filtering for genes that were detectably expressed in one or both conditions (median RPKM > 2 in either condition), we retained 10,281 genes for downstream analysis.

**Figure 1.**
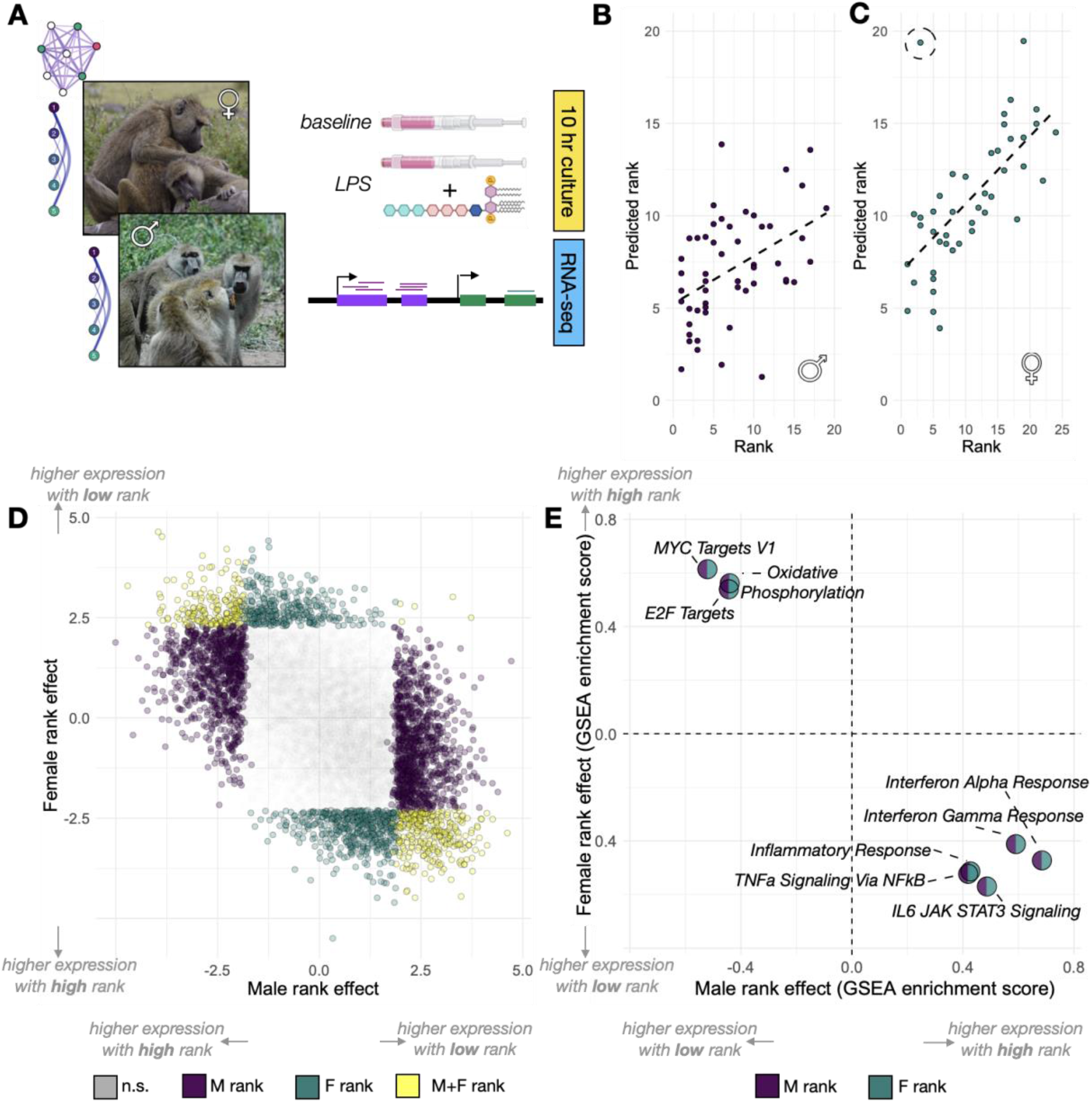
Strong, sex-specific signatures of dominance rank in white blood cell gene expression. **(A)** Study design: dominance rank (males and females) and social bond strength (females only) were evaluated for their relationship with white blood cell gene expression, generated from samples cultured for 10 hours in the absence (baseline) or presence of lipopolysaccharide (LPS). **(B-C)** A gene expression-based elastic net model accurately predicts dominance rank for male (B; Pearson’s R=0.449, p=8.46 x 10^-4^) and female (C; Pearson’s R= 0.656, p= 1.31 x 10^-6^) baboons. **(D)** The effect estimates for the rank-gene expression association are negatively correlated in males versus females (Pearson’s R=−0.544, p<10^-50^; colored dots are genes that pass a 10% false discovery rate threshold). **(E)** Gene sets enriched for higher expression in high-ranking males are enriched for lower expression in high-ranking females, and vice-versa. Enrichment in males is shown in purple; enrichment in females is shown in green. For all gene sets, enrichment score Bonferroni-corrected p-values are <0.005. Photo credits in (A): Elizabeth Archie (females) and Courtney Fitzpatrick (males). Stock images of LPS and blood draw tubes courtesy of BioRender.com.

We first investigated the signature of dominance rank separately in each sex. Consistent with our earlier work [15], we found widespread associations between male ordinal dominance rank and gene expression levels. 2,345 genes were significantly associated with male rank in baseline samples and 2,996 in LPS-stimulated samples (22.8% and 29.1% of genes tested, respectively; i.e., β_rank_ ≠ 0 in a linear mixed effects model controlling for technical covariates, age, and treatment effects, 10% FDR; Table S2. An elastic net model relating gene expression to dominance rank thus predicted male rank with moderately high accuracy (Pearson’s R=0.449, p=8.46 x 10^-4^; Fig 1B; Fig S1). In females, gene expression data were also significant predictors of dominance rank (Pearson’s R= 0.656, p= 1.31 x 10^-6^; Fig 1C; Table S3). However, the elastic net analysis for females revealed one female (AMB_2) who was high-ranking at the time of sampling (ordinal rank 3) but was predicted to be very low-ranking in both baseline and LPS samples (predicted ordinal rank 17.7 and 19.4, respectively; Fig S1). By substantially increasing our sample size and excluding AMB_2, we were able to identify female dominance rank-gene expression associations that were undetectable in previous work [15], including 1,285 rank-associated genes at baseline and 221 rank-associated genes after LPS stimulation (12.5% and 2.1% of genes tested, respectively; 10% FDR; Table S2). Because AMB_2 was a clear outlier in our sample, we report analyses excluding her in the remainder of our results; however, our comparisons are qualitatively unchanged if AMB_2 is included (Fig S2).

In both males and females, the second principal component (PC2) of the overall gene expression data was correlated with rank (males: p=1.68 x 10^-4^; females: p=0.013; note that PC1 splits baseline from LPS-stimulated cells). However, while high-ranking males tended to project onto positive values of PC2, high-ranking females tended to project onto negative values (Fig S3). Effect size estimates for individual genes were also anti-correlated by sex, such that genes that were more highly expressed in high-ranking females tended to be more lowly expressed in high-ranking males (Pearson’s R=−0.537, p<10^-50^; Fig 1D; Fig S4). As a result, genes with increased activity in high-ranking males and low-ranking females were both enriched for inflammatory and type I interferon pathways (all p_adj_<0.005; Fig 1E; Table S4) [81]. Meanwhile, genes with increased activity in low-ranking males and high-ranking females were both enriched for key metabolic and cell cycle-related pathways, including oxidative phosphorylation and *myc* signaling (all p_adj_<0.005; Table S4). Thus, the same genes and pathways were sensitive to rank dynamics in males and females, but in opposing directions. Indeed, when applying the predictive model trained for male rank to gene expression data from females, the model predictions were negatively correlated with the observed female ranks (Pearson’s R=−0.339; p=0.023), and vice-versa (correlation between female-trained model predictions and observed male rank: Pearson’s R=−0.339, p=0.014). Similarly, accessible binding sites for immune response-related transcription factors (e.g., ISL1, KLF3, defined based on increased chromatin accessibility after LPS stimulation: see SI Methods; Tables S5-S6) were over-represented near genes upregulated in high-ranking males and near genes upregulated in low-ranking females (all p < 0.05; Fig S6; Table S7).

### Distinct signatures of dominance rank and social bond strength in female baboons

To investigate whether genes that are sensitive to dominance rank (regardless of effect direction) also carry a signature of other aspects of the social environment, we next assessed the relationship between social bond strength and gene expression in the same sample. We focused exclusively on females (n=88 samples from n=44 unique individuals), using the “dyadic sociality index” (DSI), a strong predictor of lifespan in our study population that captures an annual measure of the strength of a female’s bonds with her top three female partners [45]. Female-to-female DSI is uncorrelated with dominance rank in this data set (Pearson’s R = 0.11, p = 0.468), allowing us to assess the overlap between DSI and rank associations with gene expression independently of a correlation between the predictor variables themselves. While male social bonds to females also predict male survival [45], our DSI data set for males (n=30 unique individuals) was too small to support a parallel analysis.

Controlling for dominance rank and other biological and technical sources of variance, we identified 529 DSI-associated genes (5.1% of genes tested) in female Amboseli baboons (β_DSI_ ≠ 0 in a linear mixed effects model; 10% FDR; Table S2). The vast majority of cases (522 genes, 98.6%) were specifically identified in the LPS-stimulated condition, although gene-level DSI effect sizes are correlated between conditions (Pearson’s R=0.524, p<10^-50^). Surprisingly, genes that were more highly expressed in females with strong social bonds (high DSI) also tended to be more highly expressed in low-ranking females, and vice versa, resulting in a positive correlation between the parameter estimates for rank and DSI (at baseline: Pearson’s R=0.516; in LPS-stimulated samples: R= 0.351; Fig S5; note that the positive correlation arises because low ordinal rank values reflect high rank: i.e., the top-ranked female has an ordinal rank of 1 and lower ranked females have ranks >1).

This result was counterintuitive to us because strong social bonds predict longer lifespan in Amboseli baboon females [45], but the inflammation-related pathways associated with low female rank in this population are commonly thought to be costly to health [82,83]. We therefore investigated the pathways that account for the correlation in rank and DSI effect sizes at baseline. We found that this correlation is not, in fact, driven by immune process and inflammation-related genes: social environmental effects on these genes are specific to rank, and absent for social bond strength (Fig 2A). Specifically, genes involved in the inflammatory response are highly enriched for upregulated expression in low-ranking females at baseline (p_adj_<0.005) and the majority of genes in this set exhibit a positive effect size (i.e., increased expression with lower rank: binomial test p=2.35 x 10^-12^). In contrast, there is no enrichment of inflammation-related genes for DSI (p_adj_ >.05), nor any bias in the sign of the DSI effect (p = 0.868). Consistent with these observations, accessible binding sites for TFs active in the immune response (e.g., STAT5, Smad3, STAT3) are not enriched in or near DSI-upregulated genes (all p > 0.5; Table S7). Instead, the overall correlation in rank and DSI effect sizes is driven by genes involved in cellular metabolism and cell cycle control, particularly targets of the transcription factor *myc* and genes that function in fatty acid metabolism and oxidative phosphorylation (both p_adj_<0.005; Fig 2A-B).

**Figure 2.**
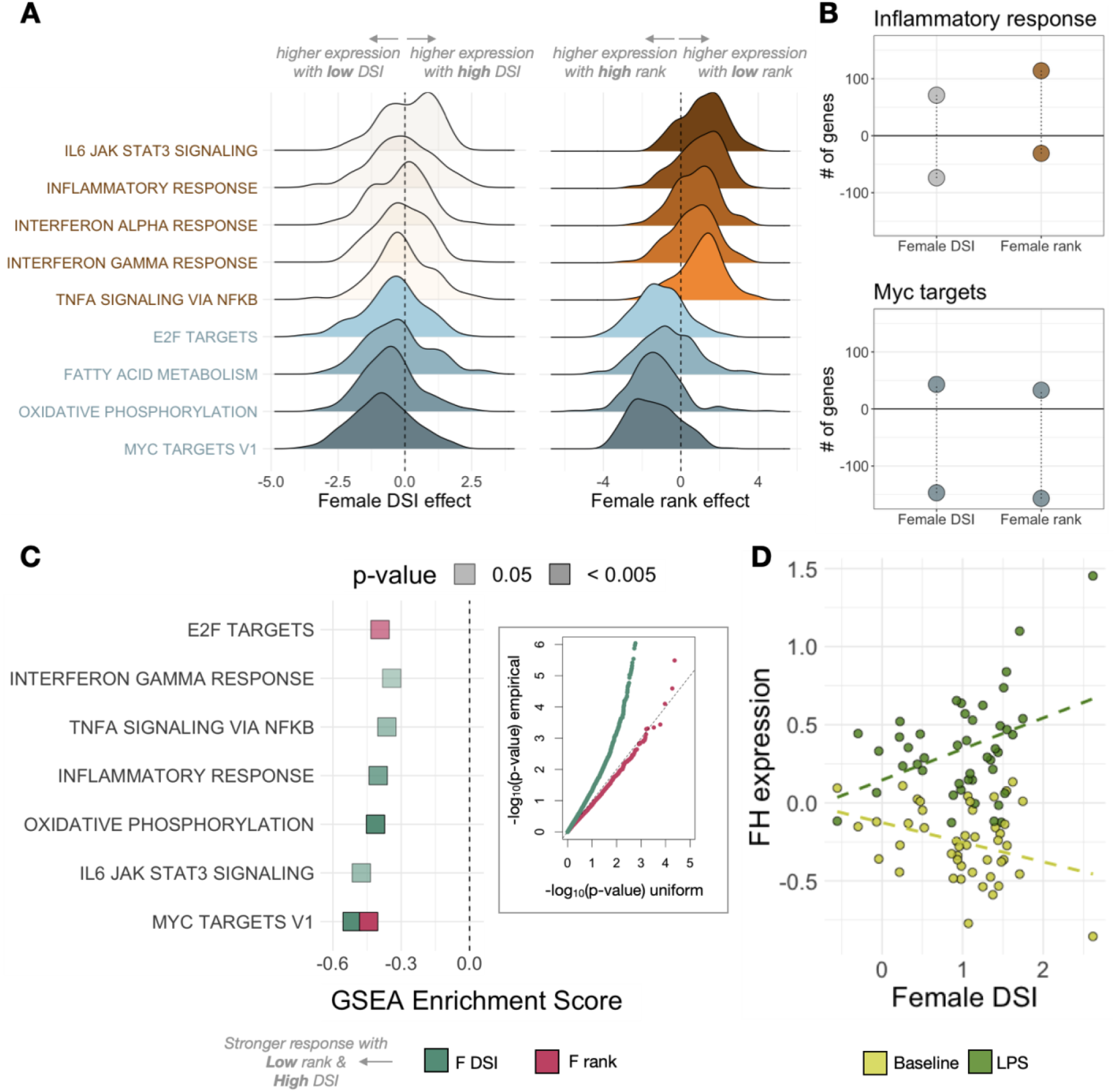
Social status and social bond strength predict distinct patterns of gene expression. **(A)** Distribution of DSI effects (left) and rank effects (right) on gene expression at baseline for genes within the MolSigDB Hallmark gene sets labeled at left. Genes within immune-related pathways (red/orange) are polarized towards higher expression in low-ranking females (positive effect sizes, because low rank is represented by high ordinal rank values). In contrast, genes in the same pathways show no pattern for association with DSI (small effect sizes centered on zero). Cellular metabolism and cell cycle-related gene sets (blue) tended to be more highly expressed with high rank (negative effect sizes) and low DSI (negative effect sizes). Translucent density plots indicate no significant bias in the direction of effects (binomial test p > 0.05). **(B)** Effect size bias for genes in the Hallmark inflammatory response and myc (v1) target gene sets, for DSI and rank respectively. **(C)** Gene set enrichment analysis results for female DSI (green) and rank (pink) effects on the foldchange response to LPS stimulation. Inset: QQ-plot of the -log10(p-value) for DSI and rank effects on the LPS response, relative to a uniform null distribution. We observe strong evidence for associations between DSI and the LPS response, but not for rank. **(D)** Example of FH, a key enzyme in the Krebs cycle that responds more strongly to LPS in high DSI females than low DSI females.

Notably, while genes involved in immune defense are not associated with DSI at baseline, a number of immune-related gene sets are significantly enriched for large DSI effects in the LPS-stimulated condition. After LPS stimulation, high social bond strength predicts higher expression of genes involved in the inflammatory response (p_adj_ = 2.0 x 10^-3^). Because these genes are not detectably associated with DSI in baseline samples, this observation suggests a potential interaction between social bond strength and the cellular environment after bacterial exposure. In support of this possibility, DSI predicts the magnitude of the *response* to LPS (i.e., the foldchange difference between LPS and baseline samples, within females) for 200 genes (10% FDR; Fig 2C; Table S8). Females with strong social bonds nearly always exhibit a more dynamic response to LPS than those with weaker social bonds (binomial test for LPS-upregulated genes: p = 1.55 x 10^-10^; binomial test for LPS-downregulated genes: p = 3.12 x 10^-12^). In contrast, because dominance rank effects are highly consistent between baseline and LPS conditions, rank does not predict the magnitude of the response to LPS (1 rank-associated gene; 10% FDR; Table S8). While many of the associations between DSI and the LPS response occur in immune pathways (Fig. 2C), females with stronger social bonds also exhibit markedly stronger responses to LPS in key cellular metabolism genes, including a key enzyme that catalyzes transitions through the Krebs cycle (*FH*: q = 0.024; Fig 2D).

### Gene expression patterns and multidimensional social advantage

To investigate the combined signatures of social status and social bond strength, we asked whether females that were relatively advantaged in both respects—and who therefore experienced advantages to both fertility and survival [44,45,75,84] — appeared physiologically distinct from other females. To do so, we binned females into four categories, corresponding to high rank/high DSI, high rank/low DSI, low rank/high DSI, and low rank/low DSI (stratified based on median rank and median DSI values in our sample). This classification reveals that, at baseline, high rank/high DSI females exhibited the lowest median expression values of genes in the Hallmark inflammatory response and IL6 signaling via JAK/STAT3 gene sets (p < 0.05 for Wilcoxon summed ranks test of high rank/high DSI females against all three other categories, for both gene sets; Fig 3). Thus, females with social capital in both dimensions—status and affiliation—present a distinct, potentially advantageous gene regulatory profile as well.

**Figure 3.**
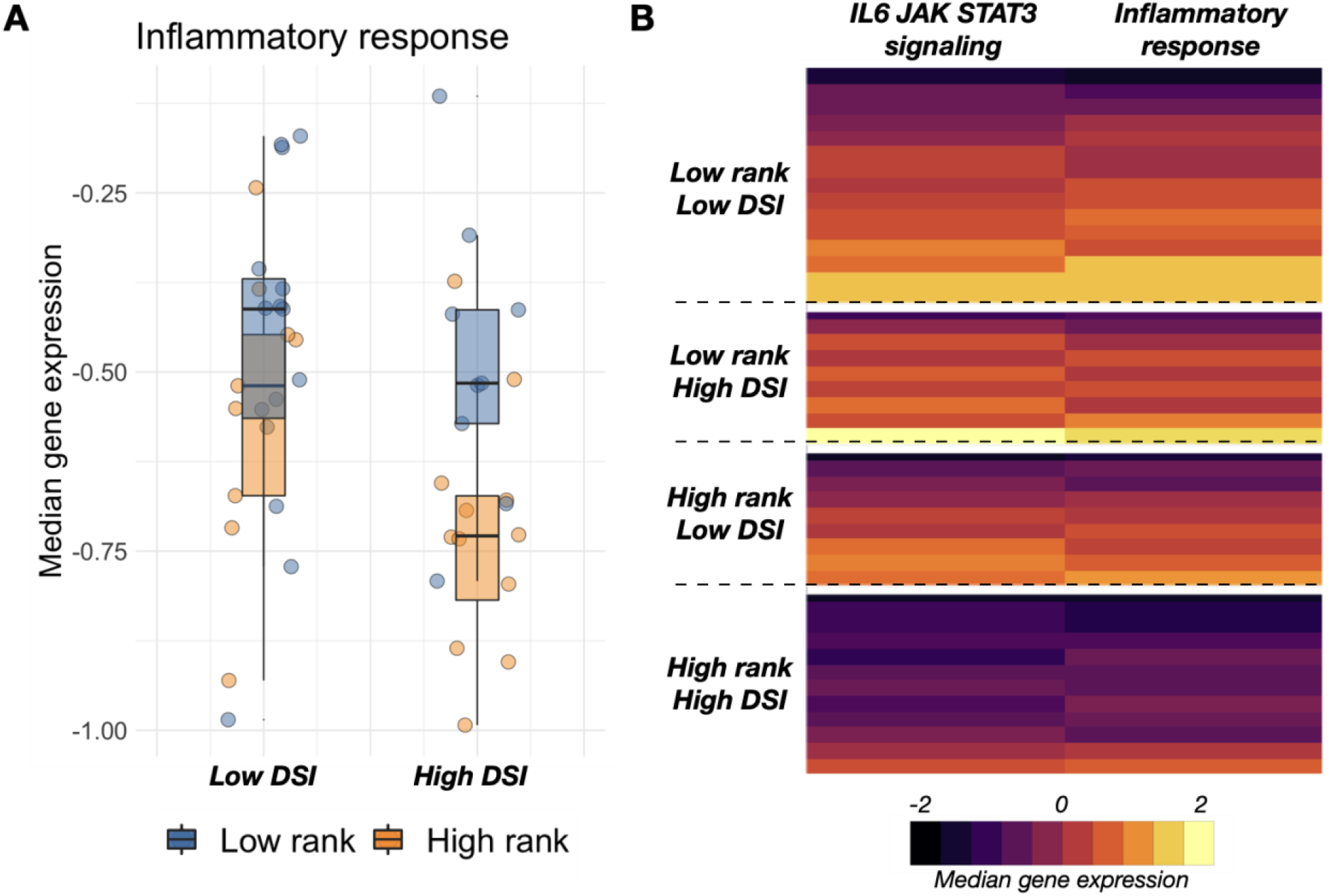
The gene expression signature of multidimensional social advantage. **(A)** Median gene expression for genes in the inflammatory response gene set illustrates that high ranking animals exhibit lower inflammation-related gene expression regardless of social bond strength (main effect of rank = −0.15, p = 0.019). There is no main effect of DSI (p = 0.166), but the difference between high- and low-ranking females is greater when high-ranking females also have strong social bonds. **(B)** Median, rescaled gene expression per individual in the Hallmark IL6 signaling via JAK/STAT3 and the inflammatory response gene sets. Each row represents a different female, with rows stratified by median dominance rank and median DSI.

## DISCUSSION

Social interactions, both affiliative and competitive, determine much about the daily experience of group-living animals. Over the life course, these experiences compound to powerfully predict health, survival, and reproductive success. Our findings reinforce that the signature of the social environment is not only observable at the whole-organism level, but also in widespread differences in gene expression. They therefore contribute to a modest but expanding body of work linking gene expression variation to social experience in natural animal populations, in both the brain and the periphery [15,70]. Together, this work generalizes extensive research on social interactions and gene regulation in laboratory models [67,85–87] to freely interacting animals in the wild. It also argues that correlations between gene expression, social status, and social integration in humans capture a broader pattern of molecular sensitivity to the social environment that predates the evolution of our own lineage [66,88].

Our findings converge with much of the previous work in humans and captive primates to indicate that innate immune defense and cellular metabolism-related pathways are closely entwined with social experience [15,57,89–92]. However, the signature of social bond strength is much more apparent after immune stimulation than at baseline, and the signature of dominance rank is substantially stronger in male versus female baboons. Thus, the functional genomic signatures of different aspects of the social environment are themselves distinct. Our results are consistent with observations that the fertility and survival consequences of male rank, female rank, and female social bond strength also differ in this population [29,44,45,75–78]. They thus call the hypothesis of a strongly conserved signature of social disadvantage into question [66]. Tests for such a signature have particularly emphasized social disadvantage-linked increases in the expression of inflammation and interferon signaling-related genes. This prediction is supported for low rank in females but not for low social bond strength—and strikingly, is directionally reversed in male baboons.

Consequently, only female social status-related differences in gene expression recapitulate the pattern reported in studies of socioeconomic status, loneliness, and social integration in humans and experimental studies of dominance rank in captive female rhesus macaques [57,89,91,93–97]. Our results suggest that low social status in female baboons may therefore be a better model for social disadvantage in humans than low social status in male baboons—perhaps because social status in male baboons is driven almost entirely by fighting ability, which is not the primary determinant of social status in modern human societies. Indeed, social environment-associated gene expression signatures in humans are often interpreted through the lens of chronic psychosocial stress [88,96,98]. While the importance of chronic stress in natural animal populations remains an open question [11], low ranking females in both this study population, wild blue monkeys, and captive rhesus macaques do exhibit higher glucocorticoid levels and/or a blunted diurnal rhythm [12,13,32,99]. Psychosocial stress may therefore be the common explanatory factor underlying conserved signatures of social adversity, when they are observed. In contrast, high rank in baboon males imposes energetic stress due to competition with other males and the demands of mate-guarding [29,100], although males may experience forms of psychosocial stress as well. And while the stability of social hierarchies and experimental work in captive primates suggests that rank precedes the gene expression patterns we observe in females, males that achieve high rank may already be physiologically distinct [15].

This explanation does not, however, account for why social bond strength does not follow the same pattern as female dominance rank. Weak social bond strength in Amboseli baboon females is also correlated with elevated glucocorticoid levels, although this effect is modest in comparison to other predictors (e.g., early life adversity [101]). If glucocorticoids are a major determinant of social environment-associated variation in immune pathway gene expression, these observations may account for why (unlike rank) we did not observe a strong signature of social bond strength in immune genes at baseline. Instead, social bond strength is most consistently linked to oxidative phosphorylation and *myc* signaling (a key regulator of cell growth, metabolism, and apoptosis). Intriguingly, *myc* activity has also been implicated in social regulation of brain gene expression in mice and as a mediator of social isolation-induced cancer susceptibility in mice and rats [64,102]. These observations suggest that social bond strength may be involved in altered energy metabolism and energetics in the baboons, as suggested in other studies of chronic and/or psychosocial stress [103].

Together, our findings emphasize substantial complexity in how the social environment is reflected at the molecular level. If we had focused only on an *a priori* subset of genes in the genome, we could have concluded that social interactions do not predict gene expression levels at all; that social status, but not social affiliation, predicts gene expression; or that social status and social affiliation generate highly similar gene expression signatures. Similarly, if we had focused only on one sex, we would have missed the shared sensitivity, but reversed directionality, of status-related pathways in males versus females. Finally, if we had only measured gene expression levels at baseline, we would have inferred that social bond strength has little relevance to immune gene regulation, when in fact it is a much better predictor of variation in the response to immune stimulation than dominance rank. While this complexity presents a challenge—additional dimensions we did not explore, including developmental, tissue, and cell type differences, are also likely to be important—it also illustrates the potential for high-dimensional genomic data to capture heterogeneity in the signature of social relationships that is impossible to infer from single measures. Indeed, our results suggest that, even in the blood, social regulation of gene expression must be the consequence of multiple upstream signaling pathways. Future studies thus have the opportunity both to test existing hypotheses about the role of glucocorticoids in social environment-associated gene regulation, and to identify alternative pathways that may also play an important role.

## Methods

### Study subjects and samples

Study subjects were 97 adult baboons (52 males; 45 females) sampled from an intensively monitored population of hybrid yellow baboons (*Papio cynocephalus*) and anubis baboons (*Papio anubis*) in the Amboseli ecosystem of southern Kenya [104,105]. Genome-wide gene expression measures were generated from blood samples collected during opportunistic dartings from 2013 – 2018. Data from samples collected in 2013 – 2016 were previously reported in [15], while the remaining 76 samples are newly reported here (Table S1). For all sampling efforts, subjects were anesthetized using Telazol-loaded darts and safely removed from their social groups for sample collection (as in [15,106,107]). Darted individuals were allowed to recover from anesthesia and released to their social group the same day.

For each study subject, we drew 1 mL of blood directly into a sterile TruCulture tube (Myriad RBM) containing cell media only (the baseline sample), and another 1 mL of blood into a second TruCulture tube containing cell media plus 0.1 ug/mL lipopolysaccharide (LPS; Fig 1A). Samples were incubated for 10 hours at 37 C. Following incubation, white blood cells were extracted and stored in RNALater at −20 C until further processing. To control for cell type composition, we also measured peripheral blood mononuclear cell type proportions for five major cell types, for each individual. To do so, we purified peripheral blood mononuclear cells (PBMCs) from blood drawn into Cell Preparation Tubes (CPT tubes; BD Biosciences) and stained the PBMCs using fluorophore-conjugated antibodies to the cell surface markers CD3, CD14, CD16, CD8, and CD20, which together differentiate classical monocytes (CD3^-^/CD14^+^/ CD16^-^), natural killer cells (CD3^-^/CD14^-^/ CD16^+^), B-cells (CD3^-^/CD20^+^), helper T-cells (CD3^+^/CD4^+^/ CD8^-^), and cytotoxic T-cells (CD3^+^/CD4^-^/ CD8^+^) [15]. PBMC composition was then profiled on a BD FacsCalibur flow cytometer and analyzed in FlowJo 10.7.1 (Table S1 with additional cell type discrimination based on cell size and granularity).

To measure chromatin accessibility, 50 mL of blood was drawn from three male anubis baboons housed at Texas Biomedical Research Institute’s Southwest National Primate Research Center into CPT tubes (BD Biosciences), spun for 30 minutes at 1800 rcf, and shipped to Duke University for PBMC isolation. 50,000 PBMCs from each individual were incubated for 10 hours at 37C and 5% CO2 in either the presence or absence of LPS (0.1 ug/mL, Invivogen ultrapure LPS from *E. coli* strain 055:B5). We then generated ATAC-seq libraries from 50,000 cells per sample (n=6 baseline and LPS-stimulated samples total from the 3 baboons; see SI Methods; [108]).

### Dominance rank and social bond strength

Sex-specific dominance ranks are assigned each month for each social group in the study population based on the outcomes of dyadic agonistic interactions observed on a near-daily basis [104,109]. Dominance rank assignments produce a hierarchy structure that minimizes the number of cases in which higher ranking individuals lose interactions to lower ranking ones [110]. To investigate rank-gene expression associations, we extracted ordinal dominance rank values concurrent with blood sample collection, which represent rank as integer values where rank 1 denotes the top-ranking individual, rank 2 denotes the second highest-ranking individual, and so on. We note that previous analyses in this and other social mammals show that alternative rank metrics sometimes confer improved predictive power [110,111]. In the Amboseli baboon population, this is especially observable in females, where proportional rank (i.e., ordinal rank scaled by group size) is more closely associated with fecal glucocorticoid levels and injury risk than ordinal rank [110]. In this data set, substituting ordinal rank for proportional rank produces highly concordant effect size estimates (R^2^ for baseline male, LPS male, baseline female, and LPS female rank effects = 0.75, 0.79, 0.88, 0.85, respectively), so we reported the results for ordinal rank for both sexes.

To measure social bond strength, we used the dyadic sociality index (DSI, as in [45,80,101]). The DSI calculates the mean grooming-based bond strength between a focal female and her top three grooming partners in the year prior to sample collection, controlling for observer effort and dyad co-residency times (see details in the Supplementary Methods). High DSI values thus correspond to strong social bonds, and low DSI values correspond to weak social bonds.

### Genomic data generation

For gene expression measurements, RNA was extracted from each sample (n=195 from n=97 unique baboons) using the Qiagen RNeasy kit, following manufacturer’s instructions (mean RIN=9.19 in a random subset of n=21 samples). We constructed indexed RNA-seq libraries using the NEBNext Ultra I or II library prep kits, followed by paired-end sequencing on an Illumina HiSeq 2500 (for samples collected from 2013 – 2016) or single-end on a HiSeq 4000 (for samples collected after 2016) to a mean depth of 17.4 million reads (± 7.7 million SD; Table S1). Trimmed reads were mapped to the *Panubis 1.0* genome (GCA_008728515.1) using the STAR 2-pass aligner [112,113]. Finally, we generated gene-level counts using *HTSeq* and the *Panubis1.0* annotation (GCF_008728515.1) [114]. We retained genes with median RPKM > 2 in the baseline samples, LPS samples, or both for downstream analysis (n=10,281 genes).

For chromatin accessibility estimates, ATAC-seq libraries were sequenced on a HiSeq 2500 to a mean depth of 40.0 million paired-end reads (± 13.7 million SD; Table S5). Trimmed reads were mapped to the *Panubis 1.0* genome using *BWA* [115]. We then combined mapped reads across samples in the same condition (baseline or LPS) and called chromatin accessibility peaks for each condition separately using *MACS2* (see Supplementary Methods; [116]).

### Gene expression analysis

To identify social environment associations with gene expression, we first normalized the gene expression data set using *voom* [117] and regressed out year of sampling (the primary source of batch effects in our data set), sequencing depth, and the first three principal components summarizing cell type composition using *limma* [118]. For each gene, we then modeled the resulting residuals as the response variable in a sex-specific linear mixed model including the fixed effects of treatment (LPS or baseline), dominance rank, DSI (for females only), age, and a random effect that controls for kinship and population structure [119]. We nested age, rank, and DSI within treatment condition to evaluate condition-specific versus shared effects. To estimate genetic covariance between individuals, which is required for the random effect estimates, we genotyped samples from the RNA-seq data using the Genome Analysis Toolkit (see Supplementary Methods; [120]). To control for multiple hypothesis testing, we calculated false discovery rates using the R package *qvalue* after verifying the empirical null was uniformly distributed [121].

To investigate how social interactions influence the *response* to LPS treatment, we calculated an equivalent to the fold-change in residual gene expression between paired LPS and baseline samples in the 44 females with both samples available. We then modeled this response using a mixed effects model, with fixed effects of age, dominance rank, and DSI, and a random effect to control for genetic relatedness/population structure. To test for enrichment of specific gene sets among rank- or DSI-associated genes, we used Gene Set Enrichment Analysis (GSEA; [122]), across the 50 Hallmark gene sets in the Molecular Signatures Database (MolSigDB; [81]). We assessed the significance of pathway enrichment scores via comparison to 10,000 random permutations of gene labels across pathways, and controlled for multiple hypothesis testing using a Bonferroni correction.

All statistical analyses in this section and below were performed in R (R version 3.6.1; [123]).

### Elastic net rank predictions

To generate predictive models for rank, we used the elastic net approach implemented in the R package *glmnet* [124]. For within-sex predictions, samples from the same treatment condition (baseline or LPS) were iteratively removed from the training set. An elastic net model was then trained using N-fold internal cross-validation on the remaining samples, and rank was predicted from the normalized gene expression data for the left-out test sample (see Supplementary Methods). To predict across sex, we trained a single model on all samples from a single treatment-sex combination, and used the model to predict rank for all samples from animals of the other sex, collected in the same treatment condition.

### Transcription factor binding motif enrichment

To investigate transcription factor binding motif (TFBM) enrichment, we focused on the 5 kb sequence upstream of rank or DSI-associated genes. We intersected these regions with areas of open chromatin called from the ATAC-seq samples, merged within treatment (e.g. the combined baseline or combined LPS samples). We then performed TFBM enrichment analysis in these regions for rank- or DSI-associated genes relative to the background set of all expressed genes using *Homer* (see Supplementary Methods) [125].

## Data accessibility

The sequencing data analyzed here have been deposited in the NCBI Short Read Archive under BioProject (PRJNA480672) for previously published data, PRJNA731520 for newly generated RNA-seq data, and PRJNA731674 for baboon PBMC ATAC-seq data. Data analysis and figure code is deposited at https://github.com/janderson94/Anderson_et_al_distinct_social_signatures.

## Acknowledgements

We thank members of the Amboseli Baboon Research Project for collecting the data presented here, especially J. Altmann for her foundational role in establishing the study population and these data sets. We also thank J. Gordon, N. Learn, and K. Pinc for managing the database, R.S. Mututua, S. Sayialel, I.L. Siodi, and J.K. Warutere for data collection in the field, and T. Wango and V. Oudu for their assistance in Nairobi. Our research was approved by the Kenya Wildlife Service (KWS), the National Commission for Science, Technology, and Innovation (NACOSTI), and the National Environmental Management Authority (NEMA) in Kenya. We also thank the University of Nairobi, the Institute of Primate Research (IPR), the National Museums of Kenya, the members of the Amboseli-Longido pastoralist communities, the Enduimet Wildlife Management Area, Ker & Downey Safaris, Air Kenya, and Safarilink for their cooperation and assistance in the field. This research was approved by the IACUC at Duke University and University of Notre Dame, and adhered to all the laws and guidelines of Kenya. For a complete set of acknowledgments of funding sources, logistical assistance, and data collection and management, please visit http://amboselibaboons.nd.edu/acknowledgements/.

