## Supplemental methods and figures for "Distinct gene regulatory signatures of dominance rank and social bond strength in wild baboons"

**Table of Contents**

1. **Supplementary Methods**
   a. Dyadic sociality index
   b. Sample processing, linear modeling, and enrichment analyses
   c. Genotype calling pipeline from RNA-sequencing data
   d. Elastic net rank predictions
   e. Measuring chromatin accessibility and TFBM enrichment analysis
2. **Supplementary Figures**
   a. Fig S1. Predicted rank for males and females in baseline and LPS samples
   b. Fig S2. Concordant effects of female rank and DSI effects when including versus excluding AMB_2
   c. Fig S3. Male and female dominance rank predict PC2 of the gene expression data in opposing directions
   d. Fig S4. Male versus female rank effects in LPS-stimulated samples
   e. Fig S5. Female rank versus DSI effects on gene expression in baseline and LPS samples
   f. Fig S6. TFBM enrichment near genes upregulated in high ranking males and low ranking females
3. **Supplementary Tables (contained within a separate Excel spreadsheet)**
   a. Table S1. Gene expression sample metadata
   b. Table S2. Results from linear mixed models for rank and DSI effects on gene expression in baseline and LPS-challenged samples (males and females)
   c. Table S3. Predicted dominance ranks from elastic net analysis
   d. Table S4. Gene set enrichment analysis results
   e. Table S5. ATAC-sequencing sample metadata
   f. Table S6. TFBM enrichment analysis results in LPS-stimulated peaks
   g. Table S7. TFBM enrichment analysis results for top 30 LPS-associated TFs nearby rank and DSI-associated genes
   h. Table S8. Results from linear mixed models for rank and DSI effects on the gene expression response to LPS challenge (females only)
4. **Supplementary References**

**Supplementary Methods**

*Dyadic sociality index*

To measure social bond strength, we used the dyadic sociality index (DSI: [1–3]), which describes social bond strength for focal adult females to their strongest social partners. DSI was calculated as a female’s average bond strength to her top three female grooming partners, in the 365 days prior to the day of sampling, controlling for observer effort. This approach is based on representative interaction sampling of grooming interactions between females, in which observers record all grooming interactions in their line of sight while moving through the group conducting random-ordered, 10-minute long focal animal samples of pre-selected individuals [4]. Because smaller groups receive more observer effort per individual and per dyad (and thus record more grooming interactions per individual or dyad), we estimated observer effort for dyad *d* in year *y* as:

$$E_{d,y}=\frac{s_{d,y}}{f_{d,y} c_{d,y}}$$

where $s_{d,y}$ is the number of focal samples taken during the dyad’s coresidence, $f_{d,y}$ is the average number of females in the group during the dyad’s co-residence, and $c_{d,y}$ is the number of days in a given year the two females in a dyad were coresident in the same social group.

DSI for each adult female dyad in each year is the z-scored residual, $\varepsilon$, from the model:

$$\log\left( R_{d,y} \right)=\beta\left( \log\left( E_{d,y} \right) \right)+\varepsilon$$

where $R_{d,y}$ is the number of grooming interactions for dyad *d* in year *y* divided by the number of days that the two individuals were coresident, and $E_{d,y}$ is observer effort.

*Sample processing, linear modeling, and enrichment analyses*

To analyze the RNA-seq data, we first trimmed the raw sequencing reads using TrimGalore (version 0.6.4; adaptor sequence “AGATCGGAAGAGC”) using default quality scores (Phred score ≥ 20) to a minimum length of 25 bp [5]). We mapped trimmed reads to the *panubis1* genome (GCA_008728515.1) using the STAR 2-pass aligner [6,7]. Splice junctions were combined across samples following the first pass alignment and used to generate an updated genome for the second round of alignment. We then generated gene-level counts using HTSeq and the NCBI *panubis1* GTF annotation (GCF_008728515.1; [8]). We focused all downstream models on genes with median RPKM >2 in either the baseline or the LPS samples (10,281 genes in total). We normalized the raw count data using *voom* with quality weights and used *limma* to remove the technical effects of sequencing depth, year of sample collection, and the first 3 principal components from the flow cytometry data [9,10]. We used the residuals from these linear models for downstream analyses.

We then modeled residual gene expression for each gene in a linear mixed effects model of the following form:

$$y_{i}\sim\mu_{i}+\beta_{i}X+g_{i}+e_{i}$$

$$g\sim MVN\left( 0,\sigma_{g}^{2}K \right)$$

$$e\sim MVN\left( 0,\sigma_{e}^{2}I \right)$$

where *y*_i_ is vector of normalized, batch-corrected gene expression levels at gene *i*, *μ_i_* is the intercept, and *X* denotes our matrix of fixed covariates and their estimated effect sizes *β_i_*. *g_i_* is a multivariate normally distributed (MVN) random effect capturing genetic non-independence between individuals (i.e., due to kinship and population structure) and *e_i_* captures residual environmental noise. *σ_g_^2^* and *σ_e_^2^* are the genetic and environmental variance components, respectively. For the kinship matrix, *K,* we used genetic covariance estimates from genotype calls generated from the RNA-seq data using the Genome Analysis Toolkit (version 4.1.3.0; see below for details; [11]).

We ran sex-specific models for each gene. For males, we incorporated the fixed effects of treatment (baseline versus LPS), age nested within treatment condition, and rank nested within treatment condition. The model for females was identical, except for the addition of DSI nested within treatment condition. For males and females separately, we controlled for multiple hypothesis testing using the false discovery rate approach implemented in the R package *qvalue* [12].

To test for enrichment of rank and DSI effects within coherent biological gene sets, we used gene set enrichment analysis [13] on the fifty MolSigDB Hallmark gene sets [14]. Specifically, we calculated the maximum absolute enrichment score for each gene set. To evaluate significance, we performed 10,000 permutations of gene IDs for each gene set and calculated the maximum absolute enrichment score for each permutation. We then constructed p-values for each gene set based on the proportion of permutation scores that were as large in magnitude, or larger, than the observed value for that set. We then performed a Bonferroni correction to control for multiple hypothesis tests.

*Genotype calling pipeline from RNA-sequencing data*

Genotype calls were generated for each sample beginning from the second-pass mapped reads from the STAR 2-pass alignment to the *Panubis* 1.0 genome [7]. We removed PCR duplicates, split N cigar reads, performed base quality recalibration, and performed joint genotyping with GATK's Haplotype Caller [15]. We then filtered genotypes with the filter: "QD < 2.0; MQ < 40.0; FS > 60.0; HaplotypeScore >13.0; MQRankSum < −12.5; and ReadPosRankSum < −8.0". We also removed genetic variants unique to a single sample. We used the sample-wise correlation matrix of the resulting genotype calls as the K matrix for modeling rank and DSI effects on gene expression.

*Elastic net rank predictions*

To predict dominance rank from gene expression levels, we used elastic net models implemented in the R package *glmnet* [16]. Specifically, we grid searched across a range of alpha values (the parameter that toggles the elastic net from LASSO-like to ridge regression-like) from 0.1 to 1, in increments of 0.1. For each alpha value, we serially removed one sample from the data set, quantile-quantile normalized the remaining gene expression values (the training set) to a standard normal distribution (first within each sample, then across samples for each gene), and built a model to predict dominance rank using N-fold internal cross-validation. The regularization parameter, lambda, was chosen to minimize the internal mean-squared error during internal cross-validation. Gene expression values for the left-out test sample were then quantile normalized to a standard normal across genes, and the value for a given gene was translated into the corresponding value observed in the training set based on the percentile in which it fell in the (normalized) training set distribution. This approach avoided the potential issue of leaking information from the training set into the test set by normalizing all the data together. For a given alpha value, the optimal model was the one that maximized R^2^ between predicted and observed dominance rank values across samples.

*Measuring chromatin accessibility and TFBM enrichment analysis*

To identify regions of open chromatin, we first trimmed ATAC-seq reads for the 3 baseline and 3 LPS samples (i.e., paired baseline and LPS-stimulated samples from each of 3 unique baboon males) using TrimGalore with default quality scores and Illumina adaptor sequence. We then mapped paired reads to the *panubis1* genome using BWA and retained uniquely mapping read pairs [17]. Following mapping and filtering, we combined read pairs within treatment (i.e., for all baseline samples and separately for all LPS-stimulated samples) across the 3 individuals, and called chromatin peaks using MACS2 [18]. We retained peaks called at a 5% FDR (“--nomodel --keep-dup all -q 0.05 -f BAMPE”) [18]. For rank or DSI-associated genes, we intersected chromatin accessibility peaks with the region 5 kb upstream of each gene. We then performed transcription factor binding motif (TFBM) enrichment analysis on accessible regions upstream of rank or DSI-associated genes. As a background set, we considered the set of accessible regions within the 5 kb interval upstream of all tested genes. We focused specifically on transcription factors important in the response to LPS, which we defined as the top 30 TFBMs enriched in ATAC-seq peaks detected in LPS-treated baboon PBMCs relative to the baseline samples (Table S6).

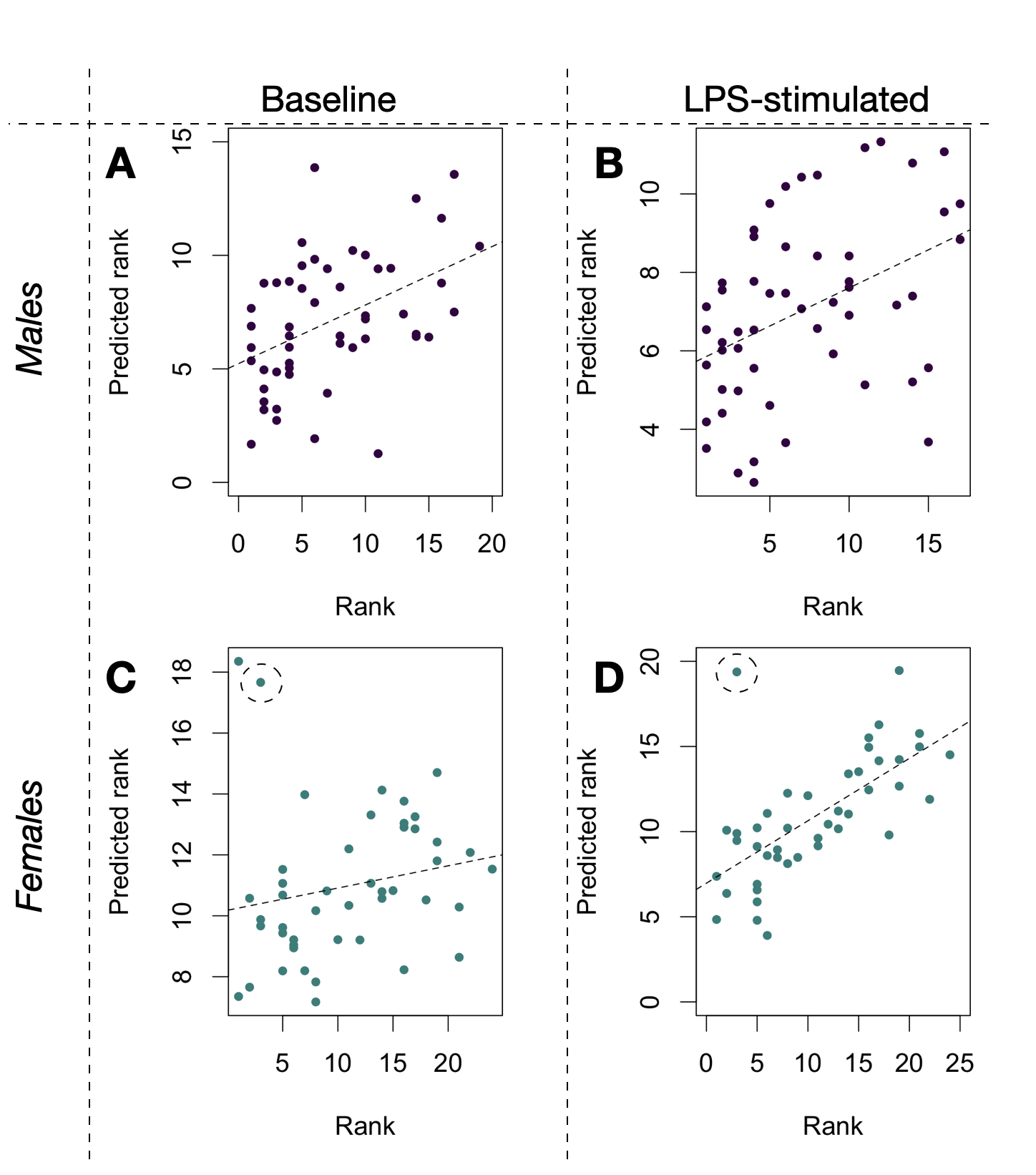

**Figure S1.** **Predicted rank for males and females in the baseline and LPS samples.** Male **(A, B)** and female (**C, D)** dominance ranks are significantly predicted by gene expression in both baseline (left column: **A, C**) and LPS-stimulated (right column: **B, D**) samples. AMB_2, the outlier female, is circled in (C) and (D). Excluding AMB_2, Pearson’s R between rank and predicted rank is R=0.449 (males at baseline, p=8.46 x 10^-4^); R=0.414 (males in the LPS condition; p = 0.002); R=0.297 (females at baseline, p = 0.047); and R=0.787 (females in the LPS condition; p = 3.95x 10^-10^).

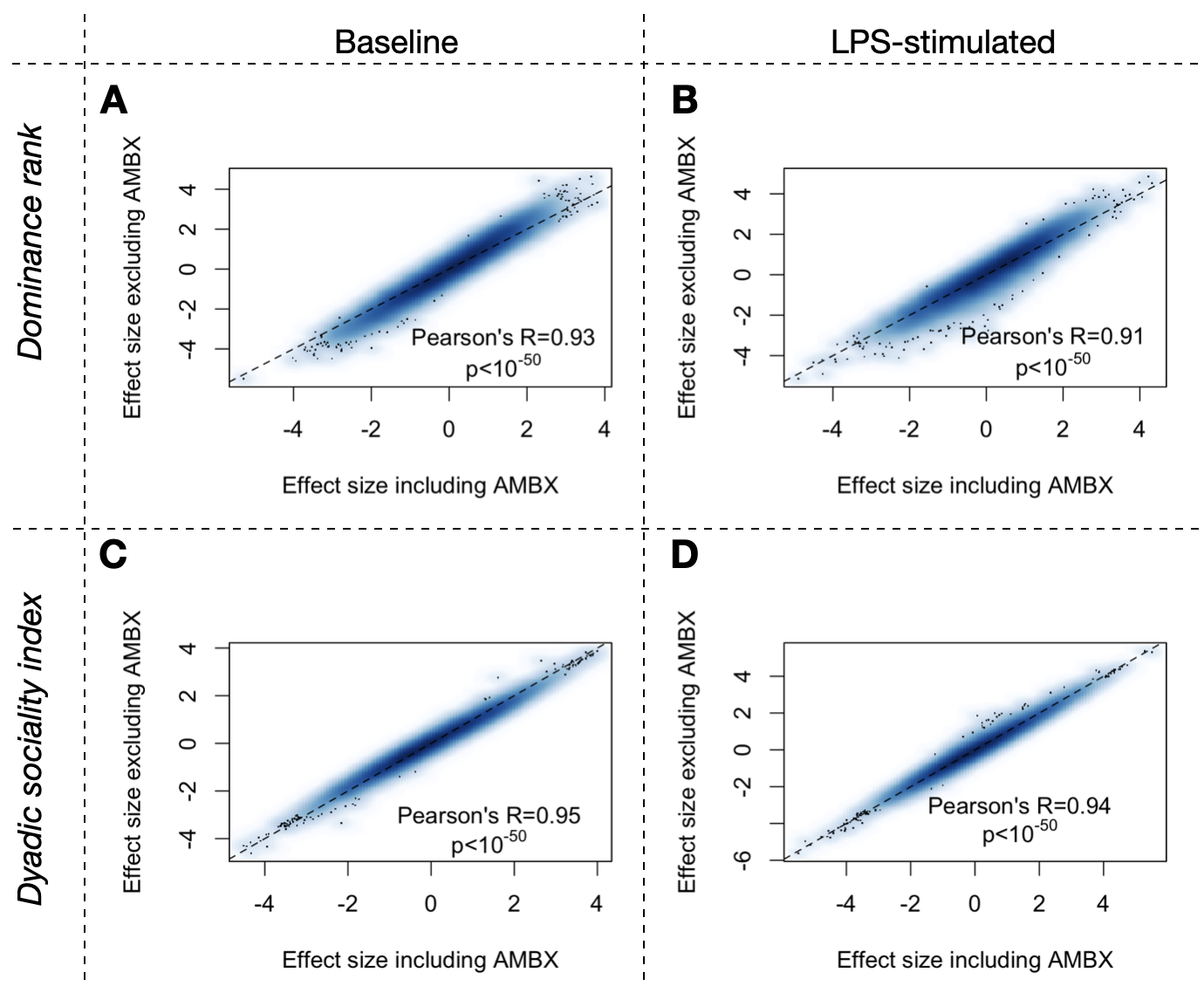

**Figure S2. Concordant effects of female rank and DSI effects when including versus excluding AMB_2.** Scatterplots comparing the effect sizes for dominance rank (top row; A-B) and the dyadic sociality index (DSI; bottom row; C-D) in baseline (left column; A,C) and LPS-stimulated (right column; B,D) samples. In all cases, effect sizes including AMB_2 and effect sizes excluding AMB_2 were highly correlated.

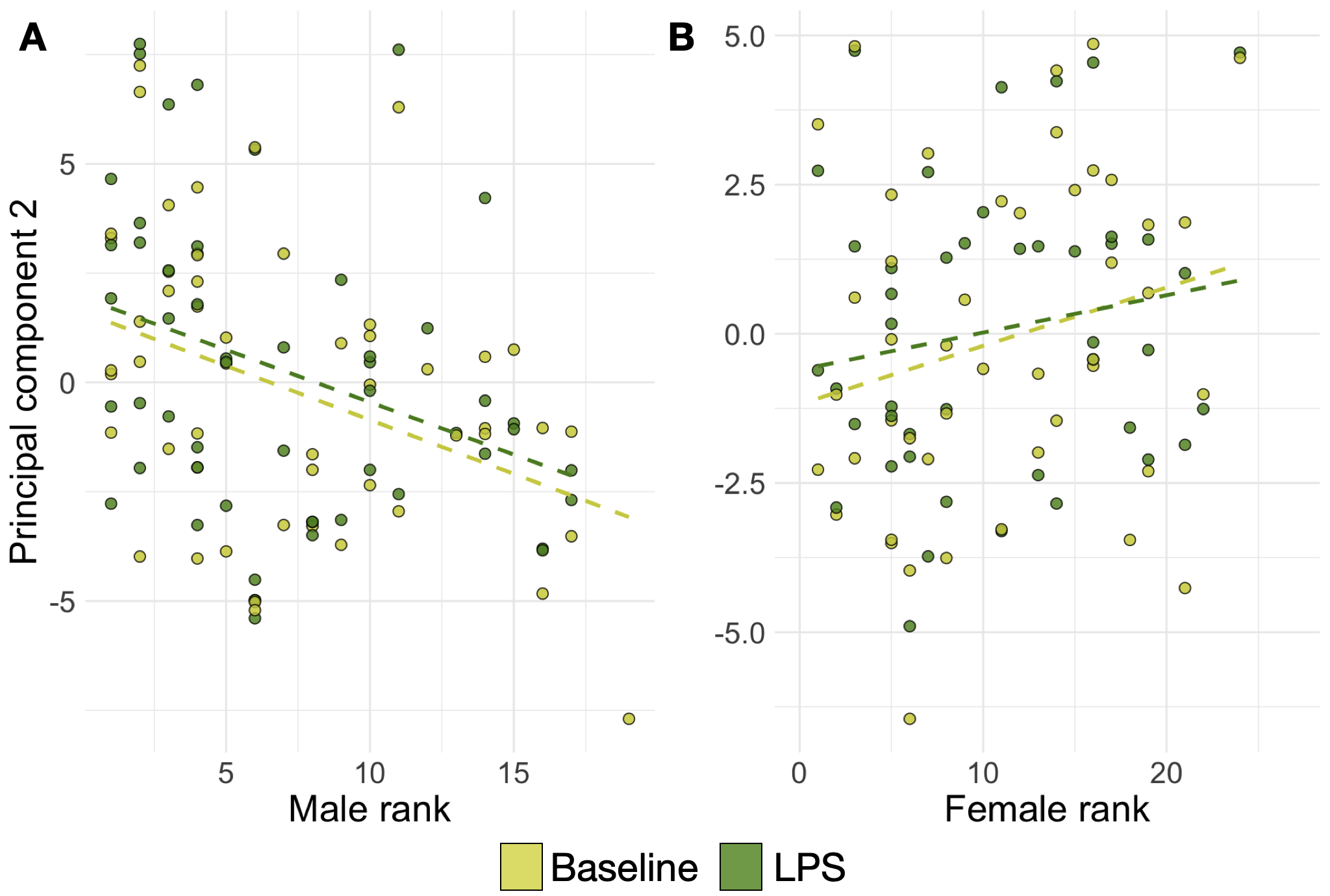

**Figure S3.** **Male and female dominance rank predict PC2 of the gene expression data in opposing directions.** Scatterplots relating dominance rank in **(A)** males, and **(B)** females to PC2 of the overall gene expression data set, including both males and females (Pearson’s R=0.359, p=1.68 x 10^-4^ in males; Pearson’s R=0.265 p=0.013 in females). The opposing direction of the rank-PC2 correlation in males and females is consistent with gene-by-gene analyses that show a negative correlation of dominance rank effect sizes between the two sexes (main text Figure 1D).

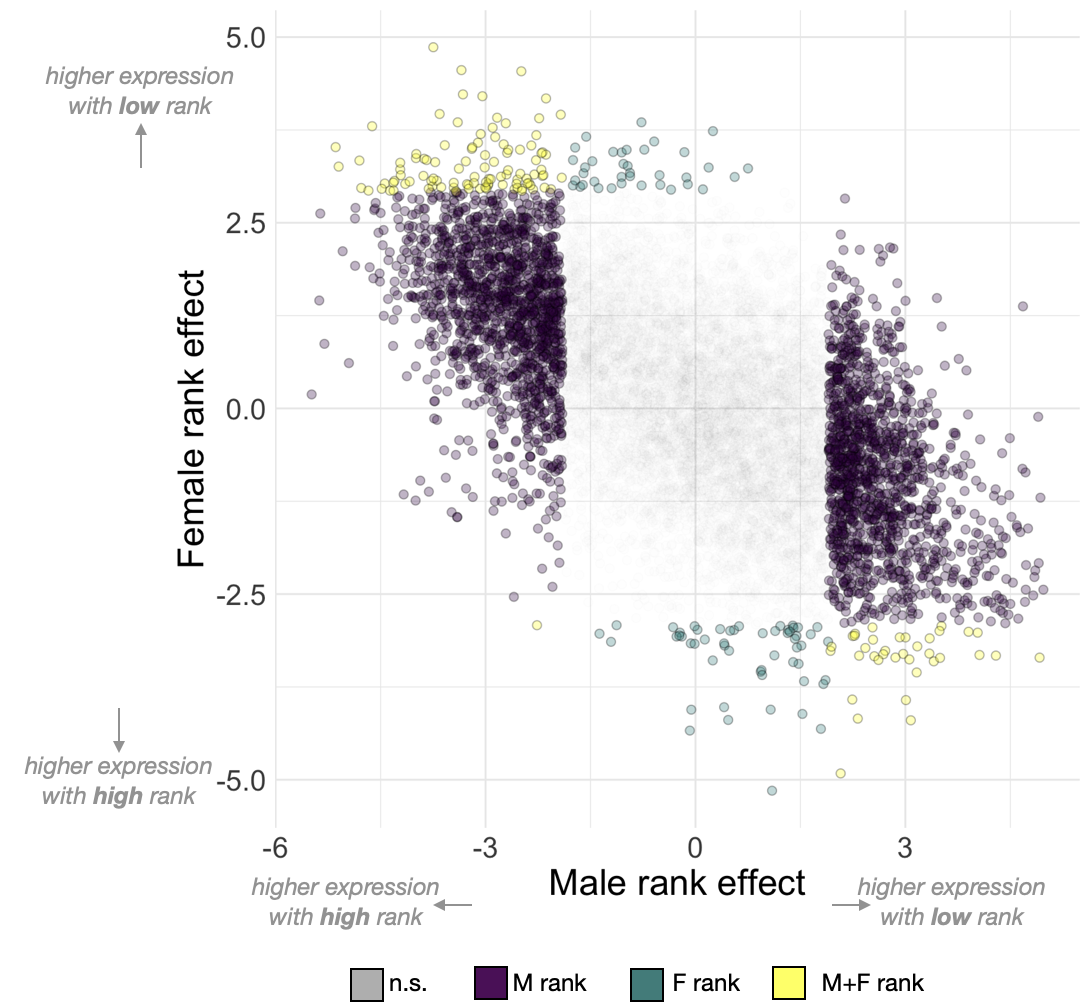

**Figure S4.** **Male versus female rank effects in LPS-stimulated samples.** The effect estimates for the rank-gene expression association are negatively correlated in males versus females (Pearson’s R=-0.577, p<10^-50^). Colors denote significance at 10% FDR. Parallel results for baseline samples are shown in main text Figure 1D.

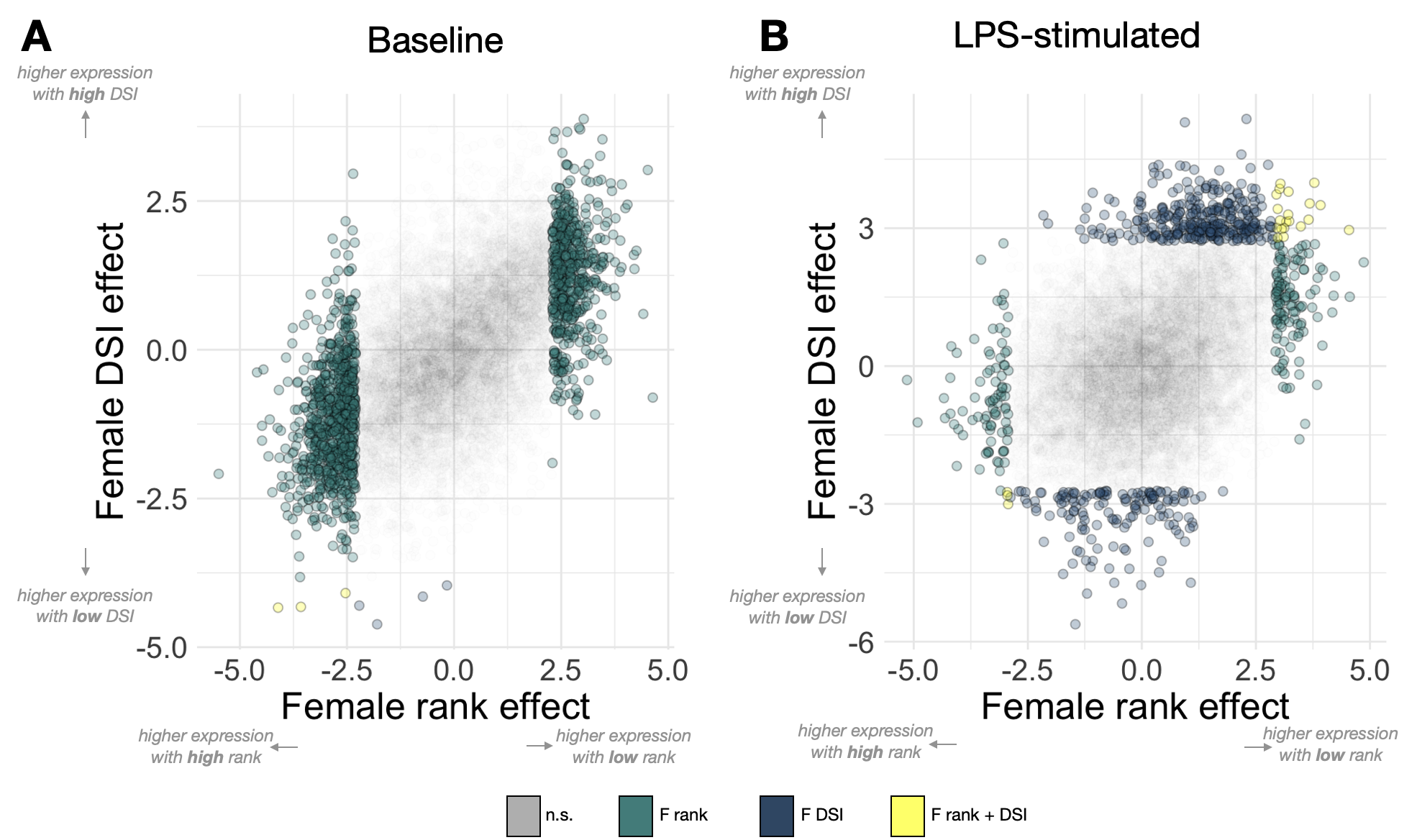

**Figure S5. Female rank versus DSI effects on gene expression in baseline and LPS samples.** The effect estimates for the rank-gene expression association and the DSI-gene expression association are positively correlated in females in both **(A)** baseline (Pearson’s R=0.551, p=<10^-50^) and **(B)** LPS-stimulated samples (Pearson’s R=0.354, p=<10^-50^). Colors denote significance at 10% FDR. The positive correlation indicates that genes that tend to be more highly expressed in low ranking females also tend to be more highly expressed in high DSI females. High expression in low rank produces positive beta values because low ordinal rank number correspond to high social status.

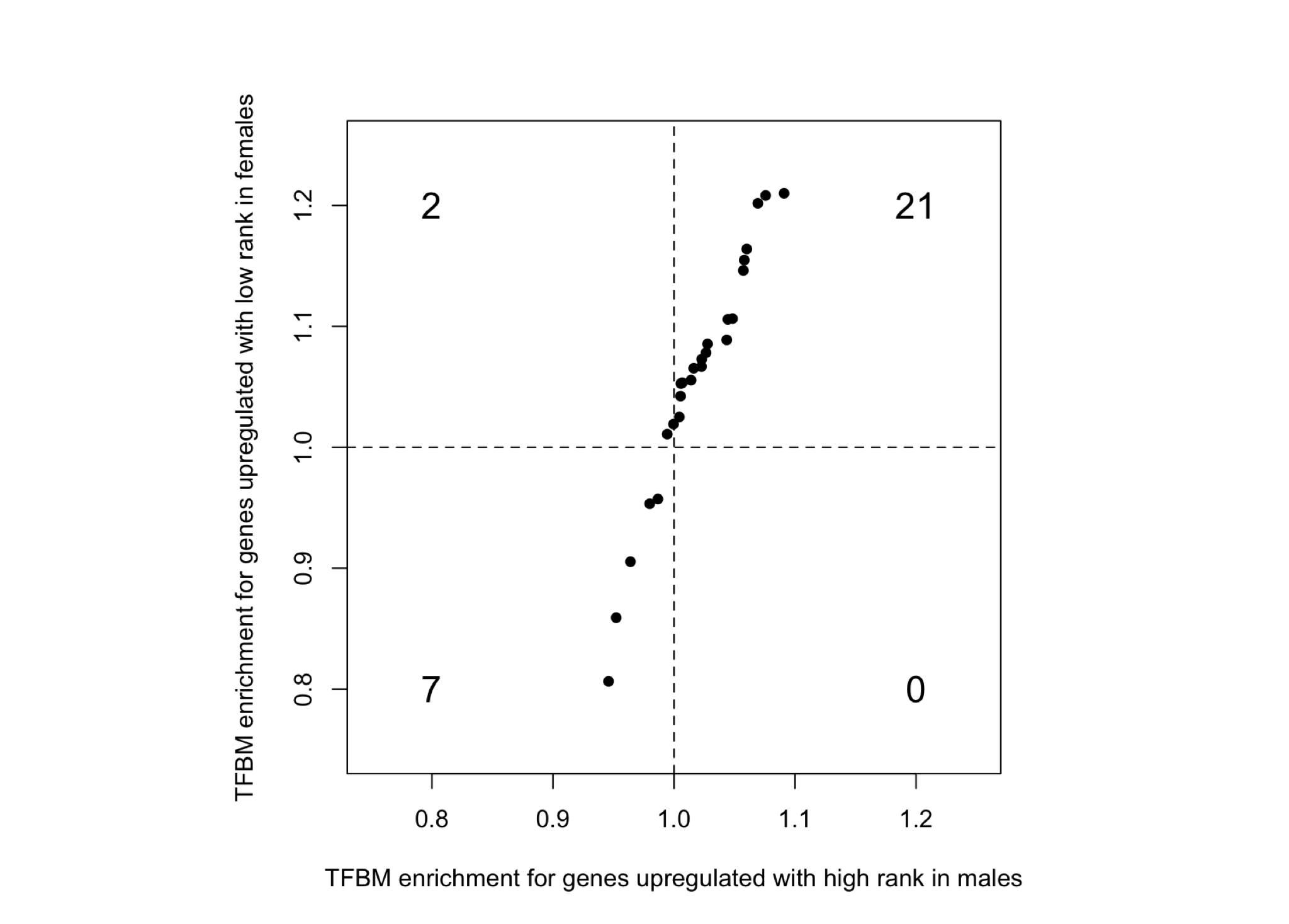

**Figure S6. TFBM enrichment near genes upregulated in high-ranking males and low-ranking females.** TFBM enrichment of the top 30 transcription factors associated with increased chromatin accessibility after LPS stimulation, in high-ranking males (x-axis) and low-ranking females (y-axis). Enrichment is expressed as the proportion of genes containing a given TFBM in open chromatin, within the 5 kb upstream of the transcription start site, relative to the background set of tested genes. Results here show enrichments for rank-gene expression associations at baseline. Numbers indicate the number of motifs in each respective quadrant.
